# Microtubule deacetylation reduces cell stiffness to allow the onset of collective cell migration *in vivo*

**DOI:** 10.1101/2021.08.12.456059

**Authors:** CL Marchant, AN Malmi-Kakkada, JA Espina, EH Barriga

## Abstract

Embryogenesis, tissue repair and cancer metastasis rely on collective cell migration (CCM). *In vitro* studies propose that migrating cells are stiffer when exposed to stiff substrates, known to allow CCM, but softer when plated in compliant non-permissive surfaces. Here, by combining *in vivo* atomic force microscopy (*i*AFM) and modelling we reveal that to collectively migrate *in vivo*, cells require to dynamically decrease their stiffness in response to the temporal stiffening of their native substrate. Moreover, molecular and mechanical perturbations of embryonic tissues uncover that this unexpected cell mechanical response is achieved by a new mechanosensitive pathway involving Piezo1-mediated microtubule deacetylation. Finally, lowering microtubule acetylation and consequently cell stiffness was sufficient to allow CCM in soft non-permissive substrates, suggesting that a fixed value of substrate stiffness is not as essential for CCM as it is reaching an optimal cell-to-substrate stiffness value. These *in vivo* insights on cell-to-substrate mechanical interplay have major implications to our re-interpretation of physiological and pathological contexts.

## Main

Cell migration is essential for a wide variety of biological processes such as embryogenesis, tissue repair and cancer metastasis^1,2^. The interaction between migrating cells and the mechanical properties of their substrates have been widely studied *in vitro*^3^ and more recently *in vivo*^4,5^. It is well established that stiffer substrates favour cell migration and *in vitro* evidence support the idea that migrating cells adjust their elastic properties to their environment^6–9^. However, recent *in silico* and *in vitro* evidence suggest that this may not be the case when cells are plated on compliant surfaces, like those observed in some *in vivo* environments^10,11^. Hence, whether and how cells that migrate in a dynamic and convoluted *in vivo* environment adjust their mechanical properties in relation to the substrate remains unclear.

To address this, we study the mechanical interplay between migrating cells and their native substrate *in vivo* by using as a model the collective migration of *Xenopus laevis* cephalic neural crest cells (NCs), a mechanosensitive embryonic cell population whose invasive ability has been likened to cancer^12^. NCs form at the border of the neural plate^13^ and it is clear that their CCM is mechanically triggered by the stiffening of the head mesoderm – the tissue that NCs use as a migratory substrate *in vivo*^4,14,15^ (**Fig. 1a**). However, whether NCs adjust its elastic properties in response to mesoderm stiffening and the molecular mechanism mediating this response remains unknown. To address this, we first measured the apparent elastic moduli (referred to here as stiffness) of wild-type mesoderm and NCs from non-migratory to migratory stages by using *in vivo* atomic force microscopy (*i*AFM) (**Fig. 1b; Supplementary Fig. 1; Methods**). Our *i*AFM measurements revealed that NC stiffness is reduced at the onset of CCM, reaching similar values to those registered in the mesoderm at this stage (**Fig. 1c**). Next, we asked whether mesoderm stiffening, which is known to trigger NCs migration^4^, mediates this unexpected decrease in cell stiffness. To test this we softened the mesoderm by using a method relying on the targeted injection of an active form of myosin phosphatase-1 (ca-Mypt1)^4,16^ (**Fig. 1d; Supplementary Fig. 2; Methods**). Targeted injection of ca-Mypt1 was sufficient to decrease mesoderm stiffness and non-autonomously block NCs migration, as we have previously shown^4^. Intriguingly, reducing mesoderm stiffness maintained NCs stiffness at similar levels to those observed in wildtype non-migratory embryos (**Fig. 1e**). These results indicate that, unlike previously proposed^6–9^, cells resting in a soft substrate are not necessarily soft and that stiff surfaces do not always induce cells stiffening, in contrast, we observed that substrate stiffening is required to reduce the elastic properties of NCs cells *in vivo*.

**Figure 1.**
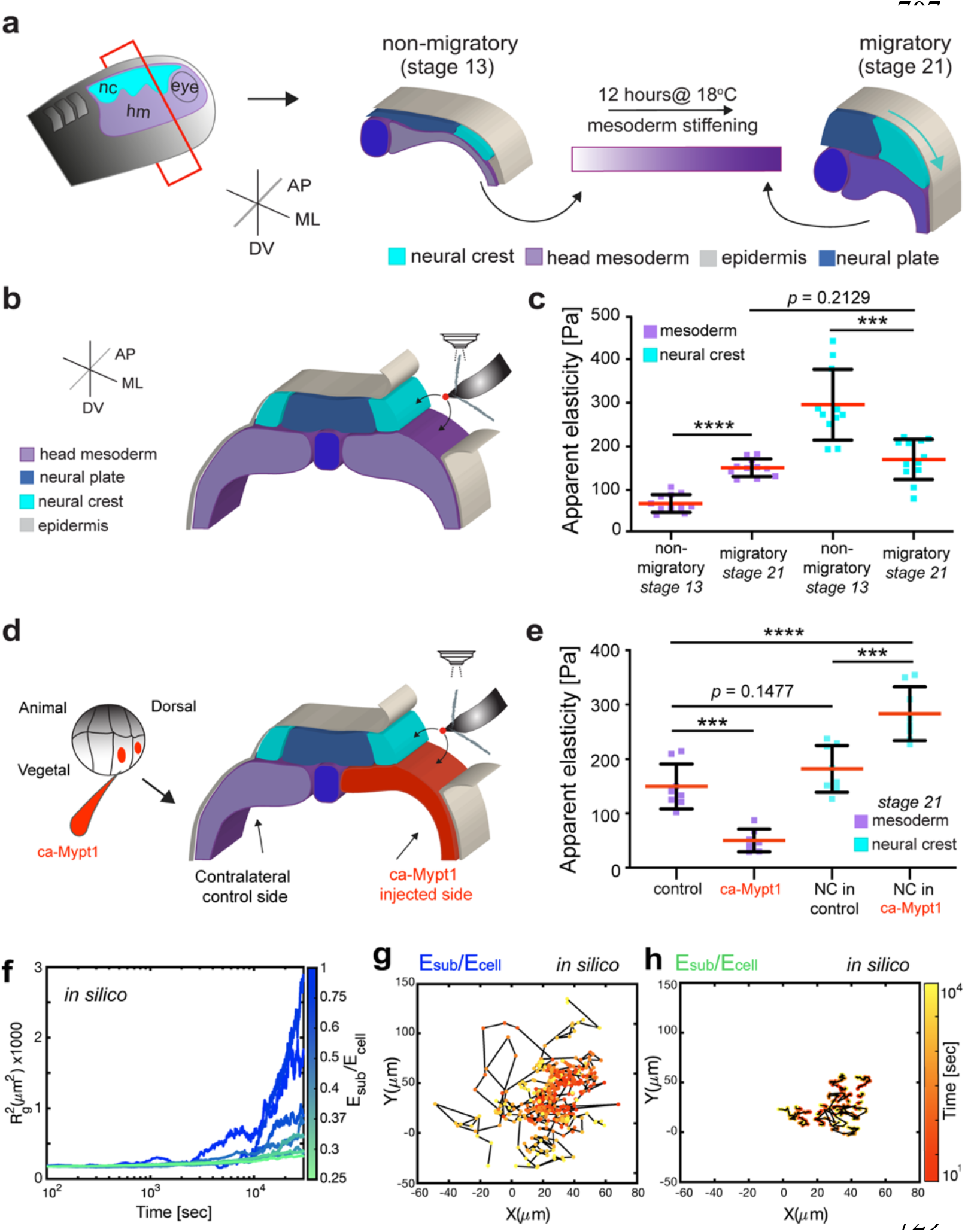
Neural crest cells reduce their stiffness upon mesoderm stiffening at the onset of CCM *in vivo*. (**a**) Diagram represents a cross-section of a *Xenopus laevis* embryos showing the development of NCs (ML, mediolateral; AP, anteroposterior; DV dorsoventral). Cephalic NC originates from ectoderm at the border of the neural plate and the onset of their CCM is triggered by stiffening of the head mesoderm, the migratory substrate of NCs. (**b,d**) Schematic showing the regions measured in wild-type or treated embryos, black arrows point to the recorded regions. (**c**) Spread of data for each condition as stated in the figure, red lines represent media, whiskers standard deviation (SD) (two-tailed t-test ****P<0.0001, ***P= 0.0004, CI= 95%, n_non-migratory mesoderm_ = 12, n_migratory mesoderm_ = 11, n_non-migratory NCs_ = 11, n_migratory NCs_ = 12 embryos (64 indentations were performed per embryo). (**e**) Spread of data for each condition as stated in the figure, red lines represent media, whiskers standard deviation (SD) (two-tailed t-test ****P<0.0001, ***P< 0.0006, CI= 95%, n_control mesoderm_ = 8, n_ca-Mypt1 mesoderm_ = 7, n_migratory NCs in control_ = 8, n_migratory NCs in ca-Mypt1_ = 8 embryos (64 indentations were performed per embryo). (**f–h**) *In-silico* results for the predicted behaviour of individual cell tracks and clusters. (**f**) 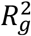 *in silico* calculations showing cell migration under the indicated conditions, line represent 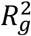 over time. (**g,h**) *In silico* cell tracks depicting individual cell trajectories (**g**) represent migration at larger ratios owing low cell stiffness values as the ones found at migratory stages and (**h**) represents poor migration at lower ratios when cell stiffness is higher, as the one reported at non-migrating stages. Results from at least 3 independent experiments or simulations.

In light of these surprising observations, we next sought to gain further insights into the impact of cell stiffness on cell migration by integrating our *in vivo* AFM data into a three-dimensional active particle computational model using the agent-based framework^17–19^ (**Supplementary Note**). Our model takes as an input the ratio of the substrate(*sub*)-to-cell(*i*) stiffness (*E_sub_*/ *E_i_*) with *E_i_* in the range of stiffness values measured for NCs from non-migratory to migratory stages (~150 to 500 Pa), and *E_sub_* was set at ~150 Pa which is the stiffness of the mesoderm at migratory stages (**Fig. 1f**). To assess the impact of (*E_sub_*/ *E_i_*) in cell movement we incorporated a self-propulsion force 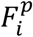 which represents the impact of cell-to-substrate mechanical interaction on the intrinsic motility of migrating cells:

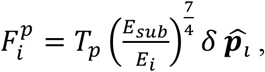

where *T_p_* is a propulsion force coefficient with units of tension (which we set to unity), *δ* the indentation of the cell into the substrate and the random polarity vector 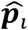 specifies the direction along which the propulsion force acts (**Supplementary Note**). Then, the impact of (*E_sub_*/ *E_i_*) in the spreading of cells within the cluster was determined through the collective variable radius of gyration squared 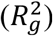, as an experimentally accessible output:

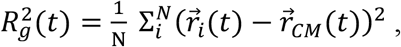

where N is the number of cells, 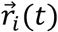 is the position of cell *i* at time *t* and 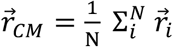 is the centre of mass of all cell positions (**Supplementary Note**). The analysis of 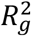 as a function of time reports the ability of the cell cluster to scatter and the tracks obtained from our simulations report on the ability of individual cell movements. Since in our experiments cell scattering is a readout of the ability of cells to migrate, we consider the radius of gyration as a measure for the efficiency of cell migration. Therefore, while low 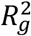 values and short cell tracks report poor migratory capabilities of the cluster and its constituent cells, larger increases in 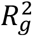 and longer tracks account for effective cell migration. Accordingly with our *in vivo* results (Fig. 1c), our simulation reported cluster deformability and cell scattering just at high values of (*E_sub_*/ *E_i_*), as shown by a rapid increases in 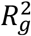 and long tracks observed in this condition (**Fig. 1f,g**). On the other hand, we found that at low values of (*E_sub_*/ *E_i_*) there is no major change in 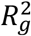 and cell migration was limited, as shown by short tracks (**Fig. 1f,h**). These *in silico* data postulate that if cells within a cluster are significantly stiffer than their substrate they should fail to migrate and *vice-versa*. In agreement to what we observed at non-migrating stages, where (*E_sub_*/ *E_i_*) was low (≤ 0.22) when compared to higher (*E_sub_*/ *E_i_*) recorded at migratory stages (~1) (**Fig.1c**).

To confirm this, we first explored what is the mechanism by which NCs adjust their elastic properties to collectively migrate. While several cytoskeletal components contribute to cell stiffness^20^, recent *in vitro* evidence proposes a central role for microtubule acetylation in tuning cell mechanics both directly and indirectly^21,22^. Given that acetylation of the lysin 40 of α-tubulin (K40-Ac) is relevant for cell motility *in vitro*^23^, an interesting possibility is that this post-translational modification could mediate the adjustment of NCs mechanics in response to mesoderm stiffening. Consequently, our *in vivo* analyses revealed that non-migratory NCs display high levels of microtubule acetylation with subsequent reduction when cells transit to a migratory stage (**Fig. 2a–c**). To confirm that this reduction in acetylation is required for the onset of CCM *in vivo* we grafted control NCs expressing wild type α-Tubulin-GFP or hyperacetylated NCs expressing an α-Tubulin mutant that mimics hyperacetylation (K40Q-GFP)^24^ into wild type host embryos (**Fig. 2d,e**; **Supplementary Fig. 3**). While control NCs grafted into wild type host embryos collectively migrated, hyperacetylated NCs displayed significant inhibition of CCM as reflected by comparing their net displacement to the control (**Fig. 2f–i**).

**Figure 2.**
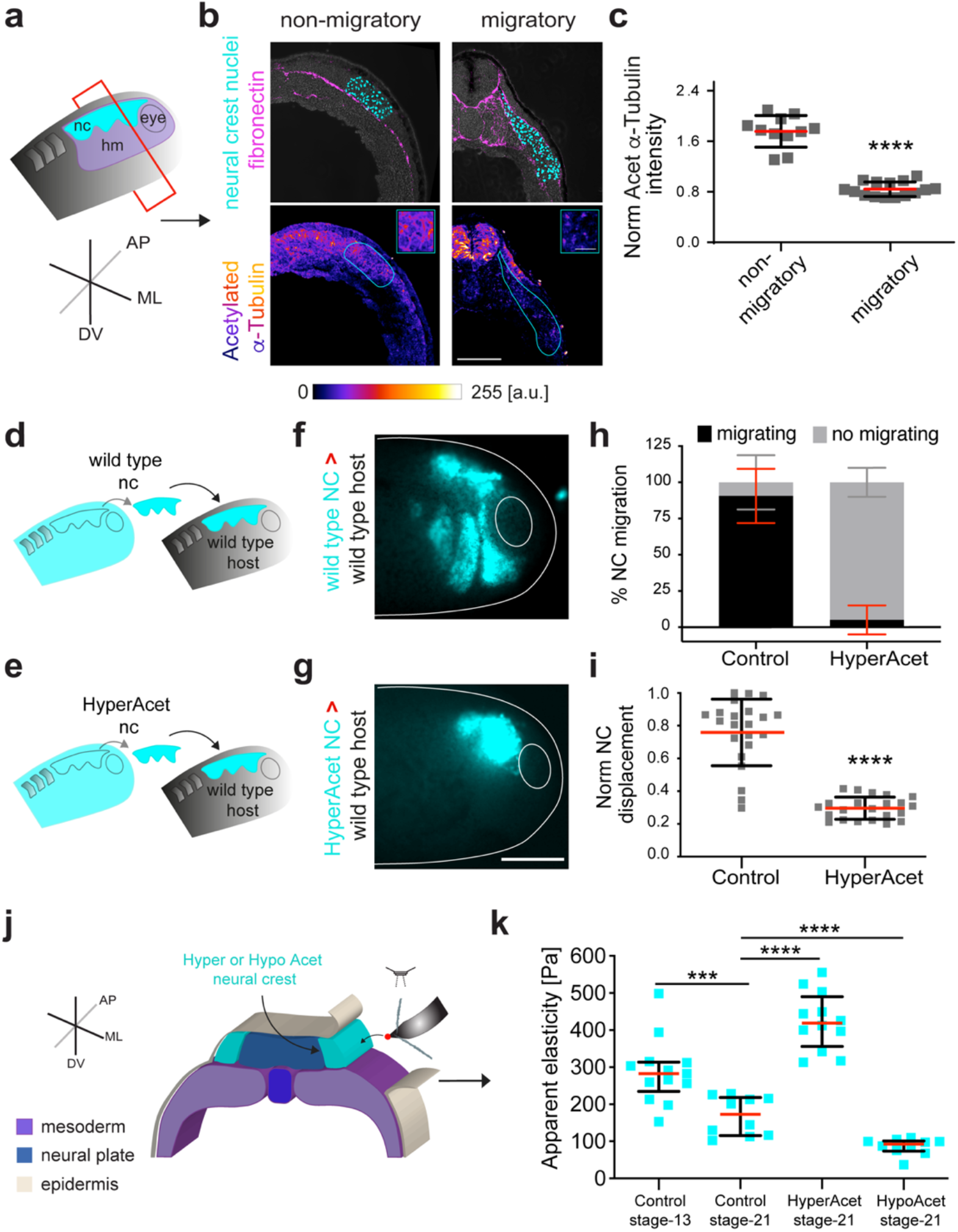
Neural crest microtubule deacetylation fine-tunes cell mechanics at the onset of CCM *in vivo*. (**a**–**c**) NCs undergo deacetylation *in vivo*. (**a**) Schematic showing the plane of sectioning. (**b**) In the upper panel, representative confocal projections of transverse cryosections showing highlighted NCs nuclei (cyan) and fibronectin (magenta) at non-migratory and migratory stages; in the lower panel, colour-coded projections of the acetylated α-Tubulin channel are shown (Scale bar 100 μm); an inset from the NCs region emphasising the signal differences between both stages is shown in the upper right corner (Scale bar 50 μm). (**c**) Normalized ratio of acetylated α-Tubulin/α-Tubulin fluorescence intensities; spread of data from the indicated conditions, red lines represent median, whiskers interquartile range (two-tailed Mann–Whitney ****P<0.0001, CI= 95%, n_non-migratory_= 17, n_migratory_= 17 embryos. (**d**–**i**) Graft experiments. (**d,f**) wildtype stage 17.5 (pre-migratory) NCs grafted into wildtype host embryos. (**e,g**) Hyperacetylated stage 17.5 NCs grafted into stage 17.5 wildtype host embryos (Scale bar for **f** and **g** 200 μm). (**h**) Percentage of embryos displaying NC migration; histograms represent media, error bars SD. (**i**) Normalised displacement of NCs along the dorso-ventral axis; red lines represent media, whiskers standard deviation (SD) (two-tailed t-test, ****P<0.0001, CI= 95. In **f** and **g**, n_control_ = 22 animals, n_hyperacetylated_ = 22 animals). (**j,k**) *In vivo* atomic force microscopy (*i*AFM) measurements. (**j**) Diagram showing the regions measured. (**k**) Spread of data for each condition stated in the figure, red lines represent median, whiskers interquartile range (two-tailed Mann–Whitney test, ****P<0.0001, ***P<0.0009, CI= 95%, n_controlNC St13_ = 13, n_ControlNC St21_ = 10, n_hypercatilated NCs St21_ = 12; n_hypoacetilated NCs St21_: 10 embryos). Panels in **b**,**f**,**g** are representative examples of at least 3 independent experiments.

Next, we analysed the impact of microtubule acetylation in the migratory behaviour of control, hyperacetylated and hypoacetylated NCs in an *ex vivo* migration assay and extracted experimental 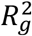 index and cell tracks for these conditions (**Methods; Supplementary Fig. 4**). Hypoacetylation was generated with an α-Tubulin mutant that mimics hypoacetylation (K40R-GFP)^24^ (**Supplementary Fig. 3**). Our *ex vivo* results confirmed that while control tracks and 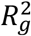 index were consistent with a migratory behaviour, hyperacetylated cells displayed shorter tracks and lower as well as constant 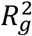 index, typical of a non-migratory behaviour (**Supplementary Fig. 4; Supplementary Video 1)**. On the other hand, the 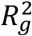 index of hypoacetylated cells displayed a rapid and overall higher increase that was consistent with large individual cell tracks, indicating that low levels of acetylation favour cell migration (**Supplementary Fig. 4; Supplementary Video 1**).

Then, we used *in vivo* AFM to quantify the impact of microtubule acetylation in cell stiffness (**Fig. 2j**). As previously observed the stiffness of wildtype NC cells showed a consistent reduction from non-migratory to migratory stages, but this trend was no longer observed in hyperacetylated NCs as this treatment yielded higher stiffness values (**Fig. 2k**). Remarkably, hypoacetylated NCs displayed extremely low stiffness values when compared to control or hyperacetylated cell stiffnesses (**Fig. 2k**). Thus, to further confirm whether the impact of microtubule hyperacetylation and hypoacetylation in CCM can be explained by their influence on cell stiffness, we integrated our observations into our theoretical framework. For this, we simulated the behaviour of cells with stiffness values recorded from control (~190 Pa), hyperacetylated (~380 Pa) and hypoacetylated (~80 Pa) NCs when migrating in a permissive substrate (**Supplementary Note**). Our simulations reproduced our experimental data with cell migration being reduced at higher hyperacetylated stiffness values and with lower hypoacetylated stiffness values yielding enhanced migration (**Supplementary Fig. 4**). These results uncover a new role for microtubule acetylation in fine-tuning NCs cell mechanics to allow the onset of collective cell migration *in vivo*.

Next, to corroborate whether microtubule deacetylation fine-tunes NCs mechanics and in turn CCM in response to mesoderm stiffening we used a controlled *ex vivo* environment which mimics the stiffness that NCs experience at non-migratory and migratory stages^4^ (**Methods**; **Supplementary Fig. 2**). In agreement with our *in vivo* observations (**Fig. 2a,b**), NCs plated on soft substrates display high levels of microtubule acetylation but these levels are drastically reduced in cells plated on stiff surfaces (**Fig. 3a,b**). Therefore, we next asked what is the molecular mechanism by which NCs sense and translate mesoderm stiffening into deacetylation to fine-tune its elastic properties and migrate. To shed light on this, we inhibited membrane mechanosensing by performing incubations with GsMTx4, an inhibitor of stretch activated channels (SACs)^25^. GsMTx4 incubation led to high levels of acetylation when compared to control cells (**Fig. 3c,d,f**), but as GsMTx4 inhibits several SACs, we next searched for specific SACs that could mediate this effect. Our RNA-seq experiments from isolated migratory NCs (**Methods**) revealed that the stretch activated channel Piezo1^26,27^ - a well-established mechanosensor^28–30^ - is significantly expressed in NCs (**Supplementary Table 1**). Thus, we tested the role of Piezo1 on microtubule acetylation by using a validated morpholino designed to knockdown *Xenopus* Piezo1 (Piezo1-MO)^28,31^. Piezo1 knockdown in the NCs led to increased acetylation levels both *ex vivo* (**Fig. 3c,e,f**) and *in vivo* (**Fig. 3g–i**), revealing a new role for Piezo1 in mediating microtubule acetylation in native contexts.

**Figure 3.**
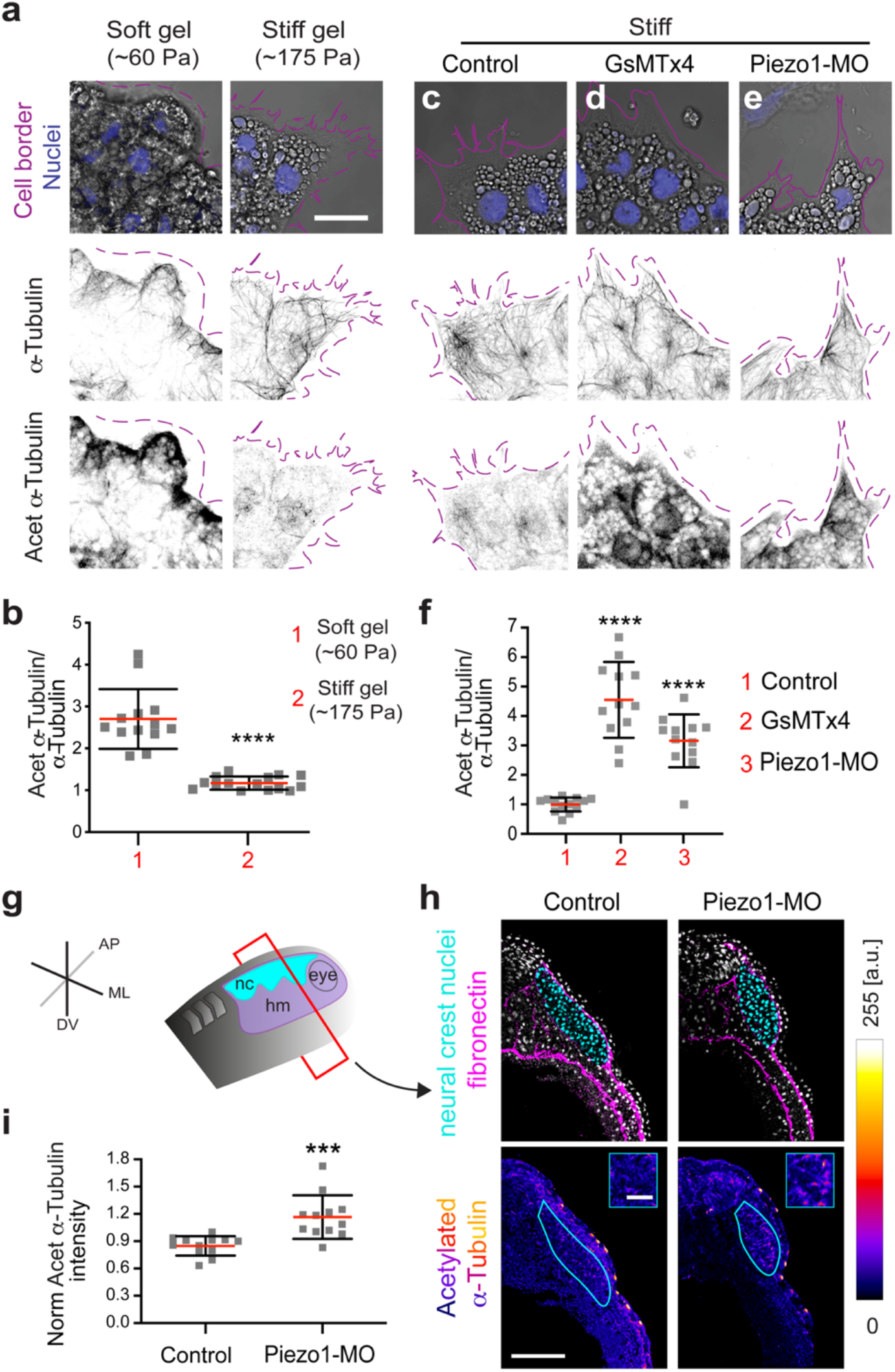
Substrate stiffness controls microtubule acetylation via Piezo1-mediated mechanosensing. (**a,b**) Immunofluorescence analysis of acetylated α-Tubulin signal of pre-migratory NCs plated on soft or stiff hydrogels. (**a**) Confocal projections, channels and conditions as indicated (Scale bar 10 μm). (**b**) Normalized ratio of acetylated α-Tubulin/α-Tubulin fluorescence intensities (two-tailed Mann–Whitney test, ****P<0.0001, n_soft_ = 13 clusters; n_stiff_ = 15 clusters). (**c–e**) Immunofluorescence analysis of acetylated α-tubulin vs α-tubulin signal in NC cells platted on stiff gels; treatments and channels as indicated in each panel. (**f**) Fluorescence intensity ratio of acetylated α-tubulin vs α-tubulin. Spread of data, red lines represent media, whiskers represent standard deviation (SD) (one-way ANOVA P<0.0001; two tailed t-test ****P<0.0001, CI= 95%, n_1–3_=37 cells). **a,c,d,e** representative examples from at least 3 independent experiments; scale bar 25 μm.

In addition to these effects in microtubule acetylation, both GsMTx4 incubation and the targeted knockdown of Piezo1 in NCs drastically impaired NCs CCM *in vivo* (**Fig. 4a–c,e,f** and *ex vivo* (**Supplementary Video 2**). Furthermore, our *ex vivo* analysis of 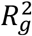 indexes revealed that the migratory ability of Piezo1 knockdown clusters and cell scattering were significantly reduced when compared to control cells, as shown by short tracks and low as well as constant 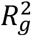 indexes (**Supplementary Fig. 5; Supplementary Video 2**) like those registered for stiff hyperacetylated cells. Since Piezo1 controls several cellular processes^32^ one possibility is that the observed effects on cell migration may be due to off-target effects. To address this, we performed an epistatic experiment in which Piezo1-MO was co-injected into NCs with a construct that mimics hypoacetylation (Piezo1-MO+HypoAcet). Strikingly, this co-injection was sufficient to rescue the effect of Piezo1 knockdown in NCs migration *in vivo* and *ex vivo*, confirming the specificity of our results (**Fig. 4d–f**; **Supplementary Fig. 5**; **Supplementary Video 2**). Next, to assess whether the defects in NCs microtubule acetylation and CCM observed upon Piezo1 knockdown are related to cell stiffness we measured the impact of Piezo1-MO in NCs stiffness by using *i*AFM. Our measurements revealed that Piezo1-MO injection in NCs was sufficient to cell-autonomously abolish the decrease of NCs stiffness that we observed in control NCs (**Fig. 4g,h**). These results indicates that Piezo1 is required to fine-tune NCs mechanics in response to mesoderm stiffening by allowing microtubule deacetylation and in turn CCM *in vivo*.

**Figure 4.**
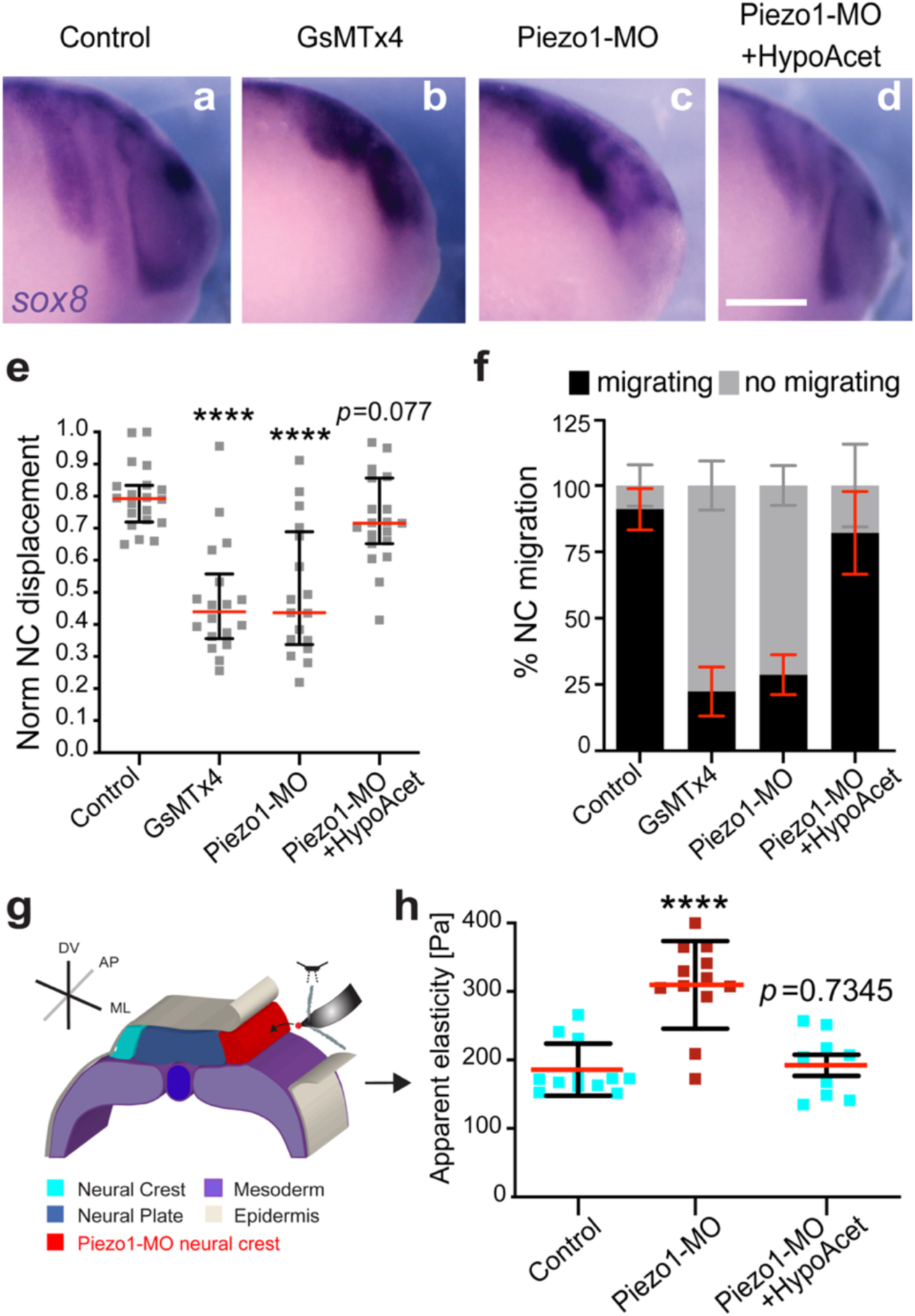
Piezo1-mediated microtubule deacetylation is required for cell relaxation and the onset of CCM *in vivo*. (**a-f**) *In situ* hybridisation analysis of NC CCM *in vivo*. (**a-d**) Lateral views of *snail2* hybridized embryos, scale bar 200 μm; treatments as indicated. (**e**) Normalised displacement of NCs along the dorso-ventral axis. Red lines show median, whiskers represent interquartile range (Kruskal-Wallis test P<0.0001, two-tailed Mann–Whitney test ****P<0.0001, CI= 95%, n_control_=20, n_GSMx4_=18, nc_Piezo1-MO_= 17, n_rescue_=19 embryos). (**f**) Percentage of embryos displaying neural crest migration histograms represent media, error bars standard deviation. Panels are representative examples from at least 3 independent experiments. (**g–h**) *In vivo* atomic force microscopy (*i*AFM) measurements. (**g**) Diagram showing the regions measured in each experiment. (**k**) Spread of data for each condition stated in the figure, red lines represent media, whiskers standard deviation (SD) (two-tailed t-test, ****P<0.0001, CI= 95%, n_Control NC_= 12, n_Piezo1-MO NC_= 12 embryos, n_Piezo1-MO+HypoAcet NC_=9 embryos).

Considering our results, our next goal was to dissect whether NCs require a threshold value of substrate stiffness to migrate or whether lowering their elastic properties to match softer substrates would be sufficient to allow CMM. Since we found that microtubule deacetylation reduces NCs stiffness to allow CCM *in vivo* we next tested whether hypoacetylation would be sufficient to allow CCM in compliant substrates, in which wildtype cells do not normally migrate^4^. As an initial approach we simulated migration of cells carrying control and hypoacetylated stiffness values when exposed to a soft substrate which resembles the stiffness of a non-migratory mesoderm (~50 Pa). Our simulations confirmed that control cells struggle to migrate on soft substrates as reflected by the shorter length of their individual tracks and almost invariable 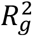 indexes (**Fig. 5a,c**). Surprisingly, individual cell tracks and 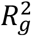 indexes of hypoacetylated NCs elicited a migratory behaviour on this soft substrates (**Fig. 5b,c**), suggesting that hypoacetylated cells could migrate in compliant surfaces. To confirm these results *in vivo*, the migration of wildtype or hypoacetylated NCs were assessed after grafting these cells into wildtype embryos or into embryos with softened mesoderm. Wildtype control NCs clusters grafted into wildtype hosts collectively migrated, but cell migration was inhibited when control clusters were grafted into softened embryos (**Fig. 5d,e,g,h**). Remarkably, hypoacetylated NCs effectively migrated by following stereotypical paths when grafted in these softened native environments (**Fig. 5f–g**). These results strongly suggest that, in this case, achieving low cell stiffness values allow reaching an optimal (*E_sub_*/ *E_cell_*) which in turn is essential for NCs to migrate on their mechanically dynamic native substrate. Consequently, we found a strong correlation between the (*E_sub_*/ *E_cell_*) calculated from our *in vivo* AFM data and the net distance that NCs migrated in embryos under all the treatments we analysed (*R* = 0.9297) (**Fig. 6a**). These results indicate that a threshold value of substrate stiffness may not be as essential for CCM as it is achieving the right cell-to-substrate stiffness ratio.

**Figure 5.**
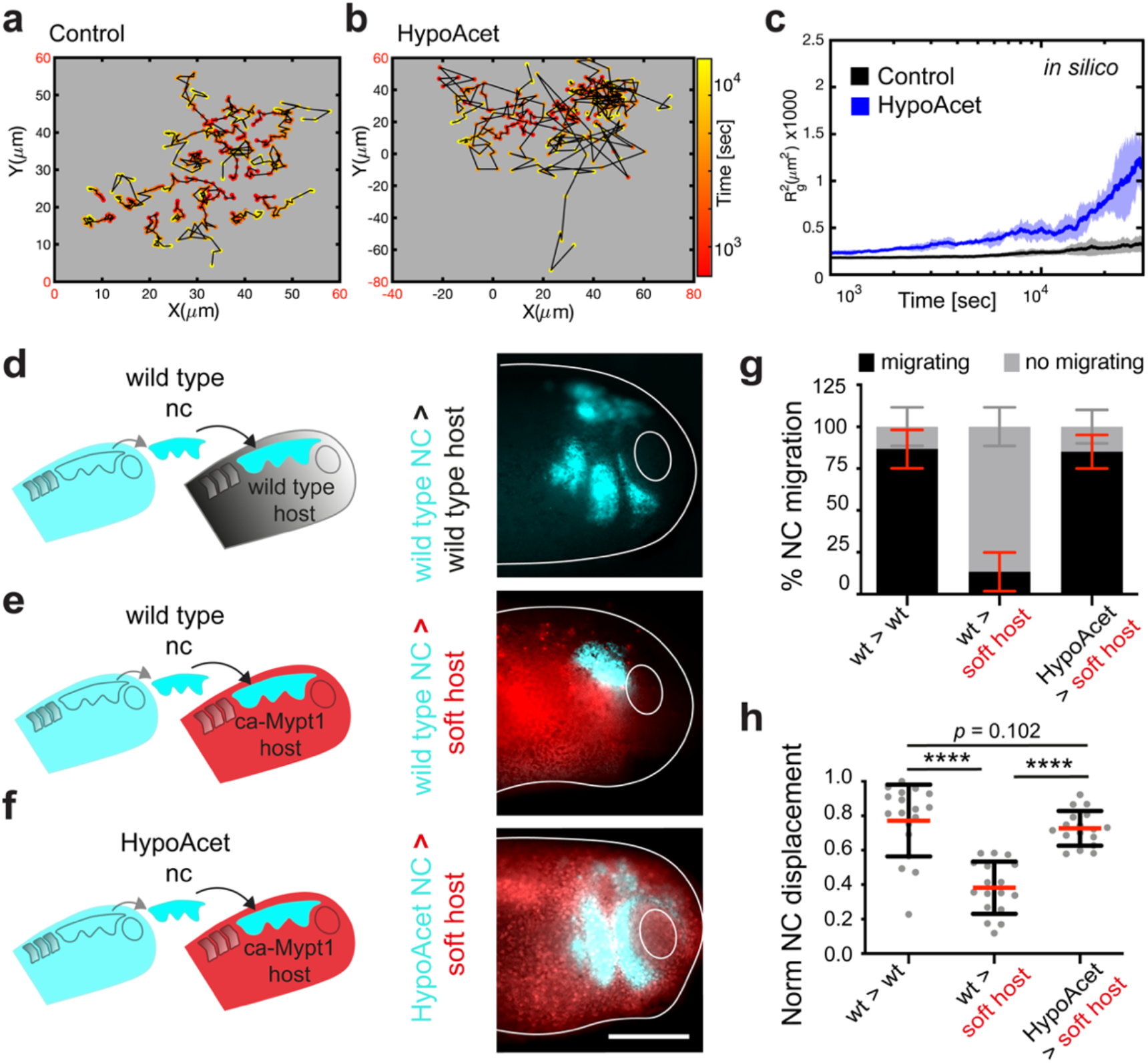
Hypoacetylation is sufficient to allow CCM in compliant native environments. (**a**–**c**) *In-silico* results for the predicted behaviour of controls and hypoacetylated cells and clusters plated on soft substrates. (**a,b**) Cell tracks depicting individual cell trajectories (note the differences in the x and y-axes scales when comparing, highlighted in red); (**c**) 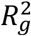 *in silico* calculations showing collective cell behaviours under the indicated conditions. (**d**–**h**) Graft experiments. (**d**) Wildtype pre-migratory (stage 17.5) NCs grafted into wildtype host embryos. (**f**) Wildtype pre-migratory NCs grafted into softened host embryos. (**f**) Hypoacetylated pre-migratory NCs grafted into softened hosts (Scale bar 200 μm). (**g**) Percentage of embryos displaying NC migration; histograms represent media, error bars SD. (**h**) Normalised displacement of NCs along the dorso-ventral axis; red lines represent media, whiskers (SD) (two-way ANOVA P<0.0001; two-tailed t-test, ****P<0.0001, CI= 95. In **g** and **h**, n_wt into wt_ = 16, n_wt into softened_ = 15, n_hypoacetylated into softened_ = 20 embryos). Scale bar 200 μm. **d–f** are representative examples from at least 3 independent experiments.

**Figure 6.**
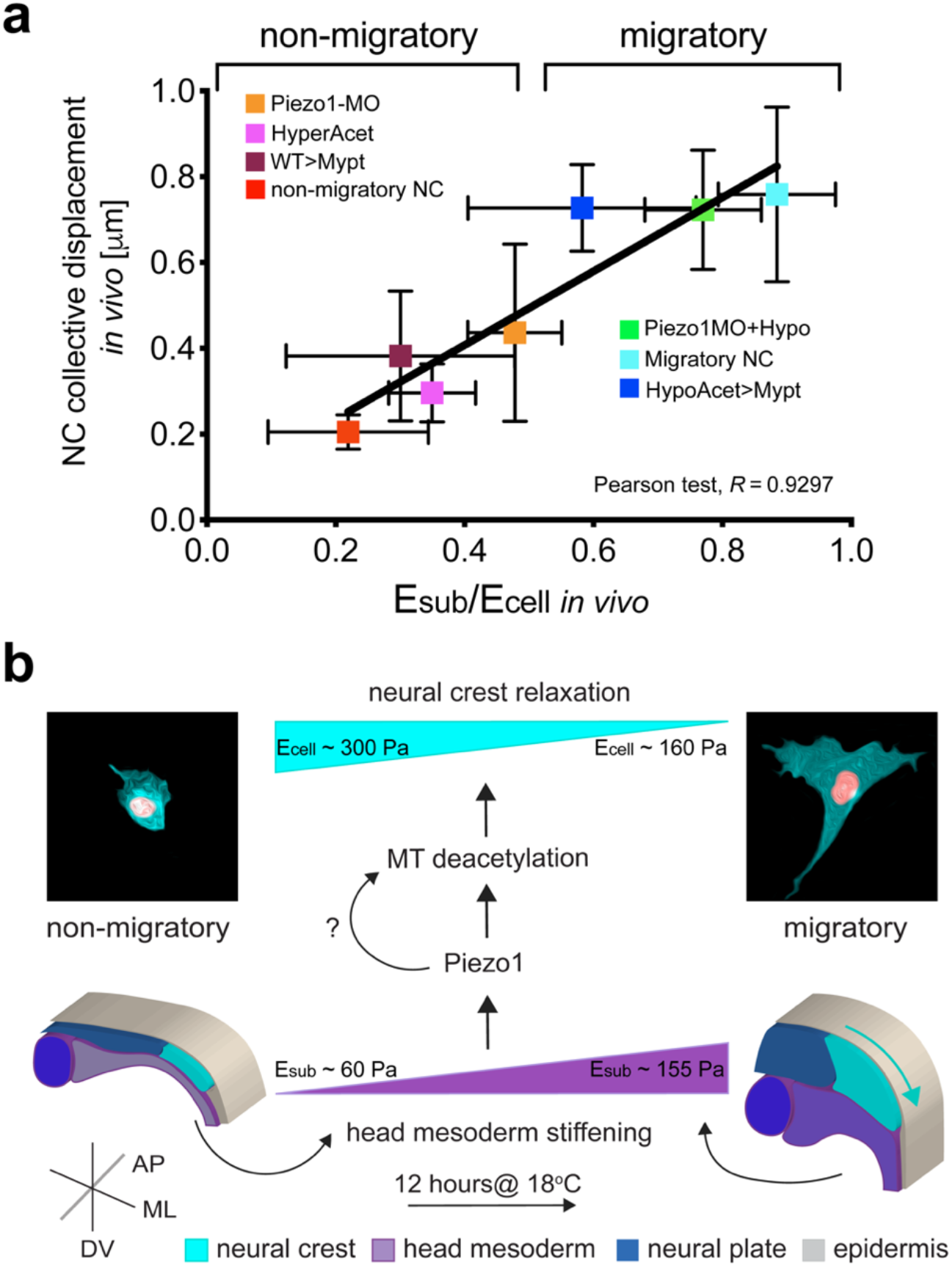
Summary of our results. (**a**) Summary of the strong correlation we found between the cell-to-substrate stiffness ratio obtained from all our *in vivo* AFM measurements and the net displacement of NCs along the embryo in under the same treatments (Pearson test, *R*= 0.9297). (**b**) Schematic providing a mechanistic overview on how the mechanomolecular feedback loop that underly onset of CCM *in vivo* operates.

Collectively, our work reveals that substrate stiffening leads to a reduction in the stiffness of migrating cells and that this unsuspected mechanical cellular response is essential for CCM *in vivo*; as it allow clusters to achieve an optimal cell-to-substrate stiffness ratio that will in turn permit cells to migrate. Mechanistically, we found that this substrate mediated reduction on cell stiffness is achieved via a new role of Piezo1 in fine-tuning the mechanics of migrating cells by mediating microtubule deacetylation (**Fig. 6b**). Thus, our data has the potential to impact our approach to several physiological and pathological processes that require CCM, such as embryogenesis, tissue repair and cancer invasion.

## Discussion

According to *in vitro* results when resting on soft substrates cells are softer than when exposed to a stiffer environment^9,33,34^. Despite this, recent evidence argue that cell and substrate stiffness are independent when cells are plated into complaint surfaces^10,11^. Our *in vivo* work reinforces this idea as we observed that NCs stiffness is higher than the stiffness recorded in the mesoderm (its substrate) prior mesoderm stiffening. In addition, our data also support an alternative or complementary scenario in which cell stiffness can be influenced by substrate stiffening, not by increasing cell stiffness as normally assumed, but by inducing cell softening and migration.

Furthermore, our work establishes a new role for microtubule acetylation in fine-tuning cell stiffness to control the onset of CCM *in vivo*. The response of microtubules acetylation to substrate mechanics show widely heterogenous behaviors *in vitro^24,35^* but whether microtubules deacetylation facilitates CCM *in vivo* was, until now, unclear. Thus, our research brings novel and physiologically relevant insights into how microtubules mediate mechanosensitive responses during CCM. Whether microtubule deacetylation directly impacts cell stiffness and/or whether it operates by controlling other downstream cellular effectors is under intense research^22^. Based on the current knowledge in the field we foresee at least three scenarios *i)* that microtubule acetylation itself could impact cell mechanics, as acetylated microtubules have been shown to be more stable and stiffer, unlike deacetylated microtubules^21,36^; *ii)* that microtubule acetylation operates by controlling the activity of GEF-H1 and with that actomyosin contractility as it has been recently proposed *in vitro*^35,37^; and *iii*) we hypothesise a combination of scenarios *i* and *ii* may emerge owing the complexity of the *in vivo* context. Regardless of whether our observations are due to a direct or indirect effect, our data position microtubule deacetylation as a key player of the cellular mechanism that modify cell mechanics to mediates tissue interaction at the onset of CCM *in vivo*.

Moreover, we reveal that Piezo1-mediated mechanosensing controls NCs’ microtubule deacetylation, cell mechanics and CCM *in vivo*. The role of Piezo1 in mediating mechanical responses has been extensively studied^26,27^. However, our results stand as the first demonstration of Piezo1 as a key regulator of microtubule acetylation and the collective migration of NCs *in vivo*. These novel results bring new challenges to the field such as gaining deeper insights in the molecular signalling by which Piezo1 controls microtubule acetylation and addressing whether this mechanosensitive pathway operates in other biological contexts. In addition, we can expect that when experiencing a soft non-migratory condition Piezo1 activity may be low. This lack of active mechanosensing could eventually explain why cells do not adjust their stiffness to the substrate in this complaint contexts, unlike in stiff substrates.

Intriguingly we also showed that inducing hypoacetylation was sufficient to allow CCM in soft native substrates. This result confirmed that lowering cell stiffness is sufficient for cells to migrate regardless the mechanical nature of their environment. Further studying these new observations can have deep implications for our understanding of processes such as cancer cell migration as recent data suggest that these cells often migrate across soft viscoelastic native tissues^38^, as the ones we report here.

Since microtubule acetylation, cell-substrate mechanics and CCM are essential for a variety of biological processes such as embryo development, tissue repair and cancer, we predict that our observations will be of general interest across the biological and physical sciences. Broadly, our data adds novel insights to the growing body of evidence arguing that mechano-molecular feedback loops, as the one described here, coordinate morphogenesis in physiology and disease^4,39–42^.

## Acknowledgements

The authors would like to thank Guillaume Charras, Ivana Pajic-Lijakovic, Marilia Henriques, Benoit Ladoux, Xin Li, Mauro Mugnai, Sumit Sinha and Dave Thirumalai for their helpful comments on the manuscript. In addition, the authors thank Joao Mata for his technical assistance along the project and for Piezo1 knockdown preliminary experiments; Dr Sofia Moreira for her help with RNA-seq experiments; and the IGC’s imaging, genomics, bioinformatics and aquatic animal facilities; Work at EHB lab receives funding from the European Research Council (ERC) under the European Union’s Horizon 2020 research and innovation programme (grant agreement No. 950254)” “EMBO IG Project Number 4765” “la Caixa Junior Leader Incoming (94978)”. EHB would like to acknowledge the support provided by Instituto Gulbenkian de Ciencia and Fundaçao Calouste Gulbenkian (start-up fund I-411133.01). AMK would like to acknowledge support provided by start-up funding from the College of Science and Mathematics at Augusta University.

## Author contributions

EHB conceived the project which evolved to its current status with major inputs from AMK and the other co-authors. CM, JAE and EHB performed experiments and measurements, and AMK developed the theoretical model and quantitative analysis tools. All the authors contributed to data analysis. EHB wrote the manuscript and prepared the figures with input from all the co-authors. JAE and CM wrote the methods section, and AMK the supplementary theory note. EHB contributed with resources for experiments and AMK for computational analyses.

## Data and code availability

Original data, AFM curves and codes that support our findings are available upon reasonable request to the corresponding author who will coordinate the delivery of the requested material. Source Data used to calculate the P values in each chart are also available.

## Methods

### *Xenopus laevis* manipulation to obtain embryos

Adult animal were maintained at 18 °C in a temperature-controlled environment and embryos were obtained as previously described^43^. Induction of ovulation was performed in adult females by injecting chorionic gonadotrophin (Intervet); after ovulation oocytes were used for *in vitro* fertilisation by mixing with a sperm solution. Embryos were staged by following established developmental tables^44^ and maintained in a temperature range between 14–18 °C. All animal experiments were approved by Ethics Committee and the Animal Welfare Body of the IGC and by the Direção Geral de Alimentação e Veterinária (DGAV). All institutional, project and personal licenses are in place.

### *In situ* hybridisation, *in vitro* riboprobes and mRNA transcriptions

*In situ* hybridizations were performed by following a previously described protocol^45^. In brief, an antisense template DNA for the neural crest marker *sox8*^46^ was generated by linearising with EcoRI (New England Biolabs). Then a digoxigenin-labeled probe against *sox8*^46^ was transcribed *in vitro* by using this linearised plasmid as a template and by following the instructions of a commercial *in vitro* Transcription System (Promega P1420). Templates for WT α-Tubulin-GFP (Addgene 56450); hyperacetylated α-Tubulin (K40Q-eGFP, Addgene 105302); and hypoacetylated -Tubulin (K40R-eGFP, Addgene 105302) were generated by PCR, using the following primers: T7-promoter containing Forward primer 5’–ggaggtctatataagcagagtaatacgactcactataggctggtttagtgaaccgtc–3’ and a Reverse primer 5’–tacgcgttaagatacattgatgagtttggacaaaccacaacta–3’. Templates for all other mRNA *in vitro* transcriptions were generated as previously described^43,47^. mRNAs for all the experiments were transcribed by following the protocols recommended in the mMESSAGE mMACHINE SP6/T7 Transcription Kits (Thermo-Fisher AM1340 or AM1334 for T7).

### Morpholino and mRNA injections

Fertilized eggs were de-jellied for 5 min with a solution containing 0.5 g of cysteine (Sigma) and 500 μl of 5N NaOH, dissolved in 25 ml of ddH_2_O. All injections were performed with pulled glass needles that were calibrated to inject 10nL upon a gas pulse of 20psi for 0.2 seconds. Depending on the type of experiment, different stages and/or blastomeres were injected (specified in each figure). For cell labelling, 250 pg of each construct (membrane GFP (mGFP) and or nuclear RFP (nRFP)) were injected per blastomere. For targeted NCs injections, embryos at eight-cell stage were injected into a dorsal and a ventral blastomeres of the animal pole with: 10ng of a morpholino designed against *Xenopus Piezo1* (Piezo1*-*MO 5’-CACAGAGGACTTGCAGTTCCATCCC-3’). This morpholino was previously validated^28^ and synthesized by GeneTools. The same strategy was used to inject the following constructs into the NC: WT α-Tubulin-GFP (Addgene 56450); hyperacetylated α-Tubulin (K40Q-eGFP, Addgene 105302); and hypoacetylated -Tubulin (K40R-eGFP, Addgene 105302)^24^.

For targeted mesoderm injections, 1ng of CA-MYPT^4^ or mGFP plasmids were injected into two dorso vegetal blastomeres at 16-cell stage, as shown in Supplementary Fig 2 and as previously described^4^.

### GsMTx4 incubations *in vivo* and *ex vivo*

For GsMTx4 incubations, embryos were incubated in a solution containing 100 μM of GsMTx4 (08GSM001, Smartox (TebuBio)) dissolved in dimethyl sulfoxide (DMSO, Thermo-Fisher). Embryo incubations were performed from stage 14 (non-migratory) until stage 22 (migratory) and immediately processed for *in situ* hybridisation. For *ex vivo* incubations, NCs explants were taken from embryos at stage 17.5 (pre-migratory) (as described below). Then NCs clusters were let to attach and spread in a fibronectin dish for 30 mins, incubated with GsMTx4 for ~2 hours and immediately processed for immunofluorescence (as described below).

### *Ex vivo* neural crest culture, dispersion assay and graft experiments

#### Neural crests dissection

This protocol has been previously described in full detail^45^. Briefly, after removing their vitellin membrane embryos were placed in a dish containing plasticine and filled with embryo media Marc’s Modified Ringer (MMR, containing CaCl_2_·2H_2_O 0.2 mM, NaCl 10 mM, MgCl_2_·6H_2_O 0.1 mM, KCl 0.2 mM, HEPES 0.5 mM with pH 7.1–7.2). Embryos were immobilised by gently holding them with plasticine and the epidermis was removed with a hair-knife tool. Then neural crest was anatomically identified and removed with the same hair-knife tool. Explants were transferred into a dish containing Danilchik’s medium (DFA; 1 mM MgSO4(7H2O), 5 mM Na2CO3, 4.5 mM KGluconate, 53 mM NaCl, 32 mM NaGluconate, 0.1% BSA and 1 mM CaCl2; after dissolving the components the pH was adjusted to 8.3 with Bicine), where they were maintained for further applications.

#### Dispersion assay

As a readout of the ability of NCs to spread and migrate we used a dispersion assay in which after dissection explants were platted into a glass bottom dish (μ-Dish, 35 mm diameter, Ibidi) that was coated with fibronectin. NCs were allowed to attach, and their migration and dispersion was recorded by time-lapse microscopy.

#### Graft experiments

Neural crest explants from wild-type or treated as well as from host embryos were removed as described in ‘*Neural crest dissection*’ Then the donor NC was carefully placed into host embryo by using a hair knife. To hold the grafted NC in place, a piece of cover-glass (0.1 mm thick) was positioned over the grafted NC. After ~1 h, the coverslip was removed, and the embryos were immediately imaged. Imaging was performed in a stereoscope as described below, by positioning the embryos on agarose dishes containing 1.2-mm lanes with 4% methyl cellulose (Sigma-Aldrich) and filled with embryo medium (MMR 0.3x). No anaesthetic was used as embryos are not motile at the analysed stages.

### Polyacrylamide hydrogels: preparation and functionalization

Polyacrilamide soft or stiff hydrogels were done as recently described by us^4^. Briefly, to generate a hydrophilic surface in which gels would attach, 76 mm × 24 mm super frost glass slides were coated with a solution made of 14:1:1 ethanol:acetic-acid:PlusONE Bind-Silene (GE Healthcare; vol:vol:vol). In parallel, gel mix solutions for either soft or stiff gels were prepared. Soft gels: 550 μL hydrochloric acid (HCL) 7.6 mM, 350 μL double-distilled water (ddH_2_O), 0.5 μL N,N,N′,N′-tetramethylethylenediamine (TEMED) (Sigma), 20 μL bis-acrylamide 2% (BioRad), 70 μL acrylamide 40% (BioRad), 5 μL of 200 nm diameter beads resuspended at 0.2 μM (Invitrogen), and 5 μL ammonium persulfate (APS) 10% (GE Health Care) (added just before use). Stiff gels: 550 μL HCL 7.6 mM, 282 μL ddH_2_O, 0.5 μL TEMED (Sigma), 25 μL bis-acrylamide 2% (BioRad), 137 μL acrylamide 40% (BioRad), 5 μL of 200 nm diameter beads resuspended at 0.2 μM (Invitrogen), and 5 μL APS 10% (GE Health Care) (added just before use). A 12-μl drop of PAA mix was placed into the hydrophilic glass slide. PAA gels were covered with a hydrophobic 13-mm diameter × 0.1 mm glass coverslips that were prepared fresh by coating them for 15 min at room temperature with PlusONE Repel-Silene ES (GE Healthcare) and dried with pressurised air. Polymerization proceeded for 40 min at room temperature in a humidifier chamber. The coverslip was carefully removed, and gels were washed 3 times for 2 min with 10 mM HEPES buffer.

### Gel Functionalization

Cell adhesion on PAA gels was generated as previously described^48^. Briefly, fibronectin was covalently linked to the soft or stiff gels by immersion of the slides into a solution containing EDC (0.2 M, (1-ethyl-3-(3-dimethylaminopropyl)carbodiimide hydrochloride), Calbiochem), NHS (0.1 M, N-hydroxysuccinimide, Sigma-Aldrich) and that were stabilised in MES buffer (0.1 M in milliQ water, pH 5.0, 2-(N-morpholino)-ethane sulfonic acid, Sigma-Aldrich) . After washing the excess of this solution with PBS, gels were incubated with 0.1 mg/ml of fibronectin for 1h 45 min at room temperature. The excess of fibronectin was removed by washing with PBS and the crosslinking-reaction was quenched by incubating the gels for 15 min with a solution containing 0.32% ethanolamine (Sigma-Aldrich) in PBS. Green Fluorescent Fibronectin (Cytoskeleton, Inc HiLyte 488) was used to determine gel functionalisation in Supplementary Fig. 2. EDC-NHS-MES aliquots can be stored at −20°C for at least 3 months.

### Cryosectioning

Embryos were fixed in PHEM-1x buffer (formaldehyde 4%; Glutaraldehyde 0.25%; Tween-20 0.1%) overnight at 4°C and then dehydrated in 100% Methanol for at least 2 h at room temperature. Then the samples were rehydrated by using a battery of methanol/PBS1x 75%-50%-25% washes, 10 min each solution and finally incubated with PBS1x. To quench auto fluorescence generated during fixation, the embryos were incubated 2 times in NaBH4 0.25% in PBS w/v for 15 min each time and finally washed with PBS1x. Then, embryos were embedded and oriented in a gelatine solution as previously described^4^. Gelatine blocks were frozen at −80 °C in pre-cooled absolute isopentane. Samples were then sectioned in 20-μm slices using a cryostat (CM-3050S, Leica) and collected in SuperFrozen Slides (VWR International). The slides were dried O/N at room temperature and processed for immunostaining, as described below.

### Immunostaining in glass, hydrogels and cryosections

Fibronectin (mAb 4H2 anti-FN, DSHB)^49^ and acetylated α-Tubulin (T6793, Sigma Aldrich)^50^ were used for immunostaining in histological cuts. To remove the excess of gelatine after cryosectioning the samples were washed twice with PBS for 15 min at 37° and blocked for 2 h with 10% normal goat serum (NGS). Antibodies were diluted at 1:500 (anti-acet-α-Tubulin) and 1:1000 (anti-fibronectin) in 10% NGS and incubated overnight at 4 °C, followed by three washes with 0.1% PBS-T (PBS, 0.1% Tween-20). Alexa-fluor (Thermo-Fisher) secondary antibodies were diluted 1:350 in 10% NGS with 1/400 DAPI added for nuclear staining. The samples were incubated in this mix overnight at 4 °C and washed three times with 0.1% PBS-T. For acetylated α-Tubulin and α-Tubulin detection *ex vivo*, explants were fixed in Buffer PHEM-1x containing (Formaldehyde 4%; Glutaraldehyde 0.25%; Tween-20 0.1%) by 10 min RT; subsequently treated with NaBH4 0.25% in PBS w/v for 10 min and washed with PBS1x; and permeabilization was done with PBS-0.1% Triton X-100 for 7 min at RT. Then the explants were blocked with 10% NGS for 30 min. The primary antibody anti-acet α-Tubulin and anti-α-Tubulin^50^ were diluted at 1:500 and 1:1000 respectively in 10% NGS and incubated overnight at 4 °C. Explants were washed three times with 0.1% PBS-Tween and incubated overnight at 4 °C with secondary antibody, diluted at 1:350 in 10% NGS. DAPI was diluted at 1:1,000 and mixed with the secondary antibodies.

Immunostaining on hydrogels proceeded as described above but the washes with agitation were replaced by rinses that were carefully performed (7 rinses each time). MOWIOL (EMD Millipore) was used as mounting medium. Images were acquired as described below and fluorescence intensity was analysed using the measurement tool from ImageJ and processed as described in statistical methods.

### Microscopy and time-lapse live-imaging

#### Time-lapse imaging

Images for dispersion assays were acquired every 5 min at 18 °C using an upright microscope Zeiss Imager Z2/Apotome.2 equipped with a motorized stage and a camera (Hamamatsu Orca flash 4.0 v2). A 10×W objective (N-Achroplan 10×/0.3 M27 (FWD=2.6mm), ZEIZZ) was used.

#### In situ hybridization imaging

All images were captured at room temperature in agarose dishes containing PBS, using a dissecting microscope (MZ FL III, Leica) equipped with a camera (DFL420, Leica) and imaging software (IM50, Leica). Magnification was 3.2×.

#### Immunofluorescence imaging

All images were acquired at room temperature using a Zeiss LSM980 system, equipped with two PMT and one GaAsPand, a 40×W objective (C-Apochromat 40x/1.1 W Corr M27, ZEIZZ). Camera, filter wheels, and shutters were controlled by Zeiss’s ZEN Blue v3.0.

### *In vivo* atomic force microscopy measurements

All AFM measurements were carried out as previously described^28^, with minor modifications. In brief, a FLEX-ANA (Nanosurf) automated AFM device, fitted with a x–y-motorized stage was used. Cantilevers coated with 5um radius colloidal spheres (CP-qp-SCONT-BSG-B-5, sQube®) were mounted on the AFM device and spring constants were calculated using the thermal noise method^51^. Cantilevers with spring constants between 0.01 and 0.03 N/m were selected. Embryos were mounted on a clay-modelling dish and prior the measurement, the epidermis was carefully dissected using a hair knife, as previously described^4^. Using the ANA software that controls the x–y-motorized stage, a region of interest (ROI) of 50×50 μm was defined in the neural crest or the mesoderm (**Supplementary Fig. 1**). Then, the following parameters were used to acquire each curve: maximum indentation force, 10 nN; approach speed, 5 μm/s; data rate, 2400 points per measurements. Force–distance curves were acquired every 6.25μm either.

### Data analysis and image treatment

#### AFM data

Apparent elastic moduli were automatically obtained by using the ANA software provided in our FLEX-ANA AFM setup. To do this, force–distance curves were fitted to a Hertz model for a spherical indenter.

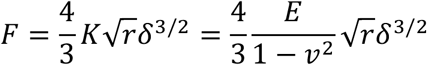

with applied force *F*, Young’s modulus *E*, Poisson’s ratio *ν*, indenter radius *r*, indentation depth *δ*, and apparent elastic modulus *K* = *E*/(1 – *ν*^2^), which is referred to as “stiffness” in the text and as ‘apparent elasticity’ in the y-axis of each chart. In all our experiment, 64 indentations were performed per embryo and obtained force–distance curves were selected as previously described^4^. Then the median of each embryo was calculated and processed for further statistical analyses.

#### *In vivo* analysis of neural crest migration

For *in situ* hybridization and grafted embryos, the length of the neural crest was obtained and normalized against the total dorso-ventral length of the embryo. Lengths were obtained using the built-in measurement tool from ImageJ and further analysed as described in Statistical analysis.

#### *Ex vivo* analysis of neural crest migration

*Dispersion assays*, an ImageJ-based Delaunay-triangulation plugin was used to calculate the distance between neighbor cells. Data were further analysed as described in the Statistical analysis section. *Squared radius of gyration* 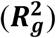, cell tracks were obtained with ImageJ Manual Tracking plugin and the obtained tracks were processed for 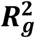 analyses. 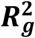 calculation and plots were done by using custom made MATLAB codes (available upon request).

### Image treatment

*z*-stacks, maximum projections and time-lapse movies were created using ImageJ software. Adjustment of display map levels, re-sizing, and addition of scale bars and pseudo colour were applied with ImageJ and/or Adobe Photoshop.

### Statistical analysis

To determine the size of the sample we follow published studies and no statistical method was done. No randomization of the experiments was performed. Owing to the nature of our experiments, only viable embryos and cell clusters were considered for analysis. Moreover, mis-injection was not included for *in situ* hybridization analysis (proper injection was assessed as described above) meaning that the authors were not blinded to allocation while performing and/or analysing the experiments. For any of the mentioned cases, after selections, all parameters were measured at random.

Each experiment was repeated at least three times. Every set of data was tested for normality test using the, d’Agostino–Pearson and/or Shapiro–Wilk test in Prism7 (GraphPad). For paired comparisons, significances were calculated Prism7 with a Student’s *t*-test (two-tailed, unequal variances) when the distributions proved to be normal. If a data set would not pass the normality tests the significances were calculated with Mann-Whitney (two-tailed, unequal variances). For multiple comparison of data with normal distribution unpaired one-way analysis of variance (ANOVA) with Bonferroni test correction was performed; while non-normal distribution data sets were analysed with Klustal-Wallis corrected with Dunn’s test. Individual comparisons were calculated only when multiple comparisons showed *P* > 0.05 and significances in these cases were calculated in Prism7 as described for paired comparisons. Confidence interval in all experiments was 95% and as detailed description of statistical parameters it was included in all figure captions.

