## Supplementary Information for "Microtubule deacetylation reduces cell stiffness to allow the onset of collective cell migration *in vivo*"

**Supplementary Figures:**

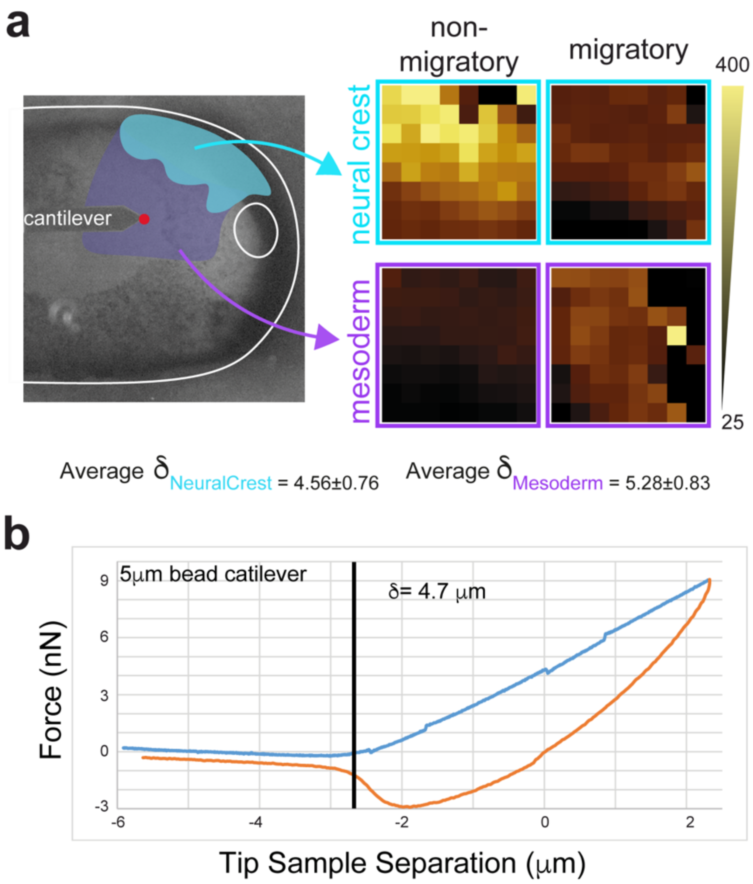


**Supplementary Figure 1. *in vivo* Atomic Force Microscopy experiments.** (**a**) Image of the AFM cantilever position relative to the neural crest (cyan) and mesoderm (magenta). A heat map showing 64 measurements acquired from an 8x8 grid with 6.25um of separation between each measurement is shown; this grid depicts the mechanical heterogeneity found in the tissues we measured. All measurements were recorded by using a 10 □m bead attached to the tip of our cantilevers. Each data point in our charts presented in the main figures represent an embryo from which the median resulting from our 8x8 grids was calculated. (**b**) Representative example of a force-distance curve obtained using cantilevers coated with 10-µm beads. Average indentation depth across all our measurements with its respective standard deviation is also shown.

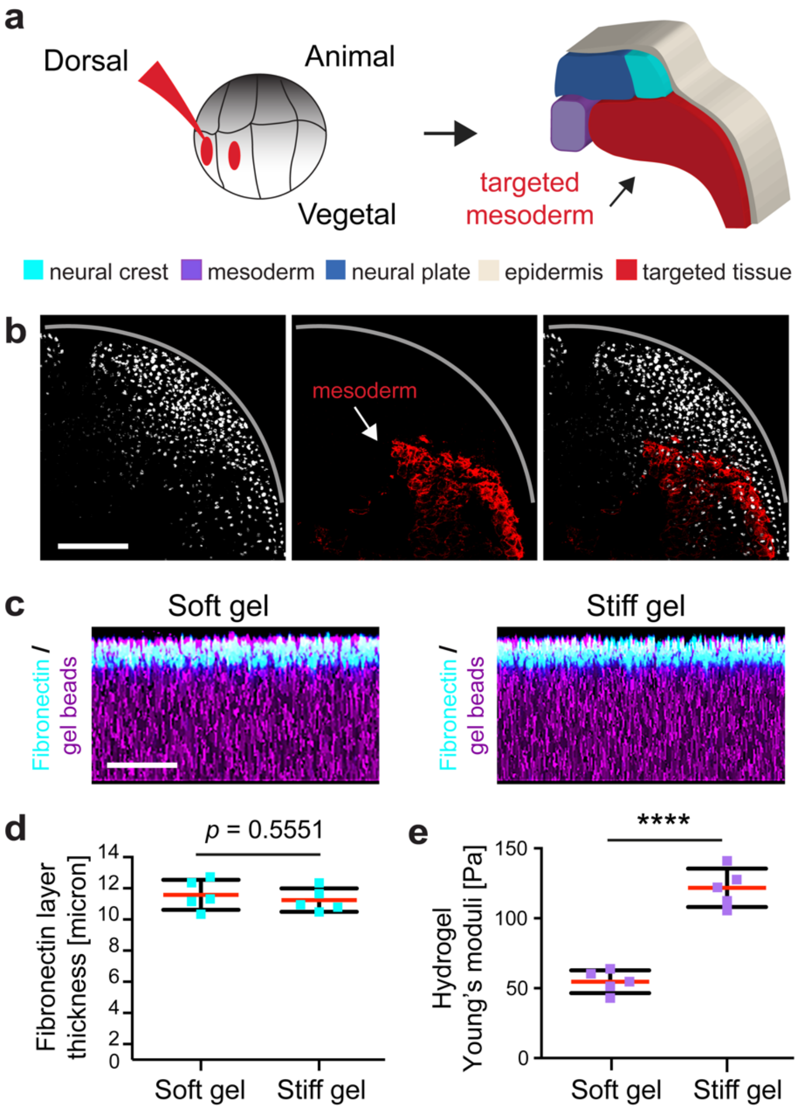


**Supplementary Figure 2. Targeted injections into the head mesoderm.** (**a**) Schematic displaying how to target the mesoderm. (**b**) Confocal projections showing the result of our targeted injections into the mesoderm (red). Scale bar, 100 µm. (**c**–**e**) Characterisation of the *ex vivo* system that reproduces the stiffness values that neural crest cells experience at non- and migratory stages. (**c**) Orthogonal view of a confocal projection of soft and stiff hydrogels. Images in (**c**) are representative examples from at least 3 independent experiments; scale bar, 50 µm. (**d**), Chart showing that the layer of Fibronectin has similar thickness in soft and stiff gels; spread of data is shown (data points represent the average obtained from each gel and 5 measurements were taken from each gel), red line represents mean and whiskers show standard deviation (s.d.); n = 5 gels; two-tailed t-test, CI = 95%. (**e**) AFM measurements obtained from soft and stiff hydrogels; spread of data is shown (each data point represents the average of a gel, and 64 measurements were taken per gel), red lines show media and whiskers s.d.; n = 5 gels; two-tailed t-test, ****P < 0.0001, CI = 95%. Scale bars, 50 µm.

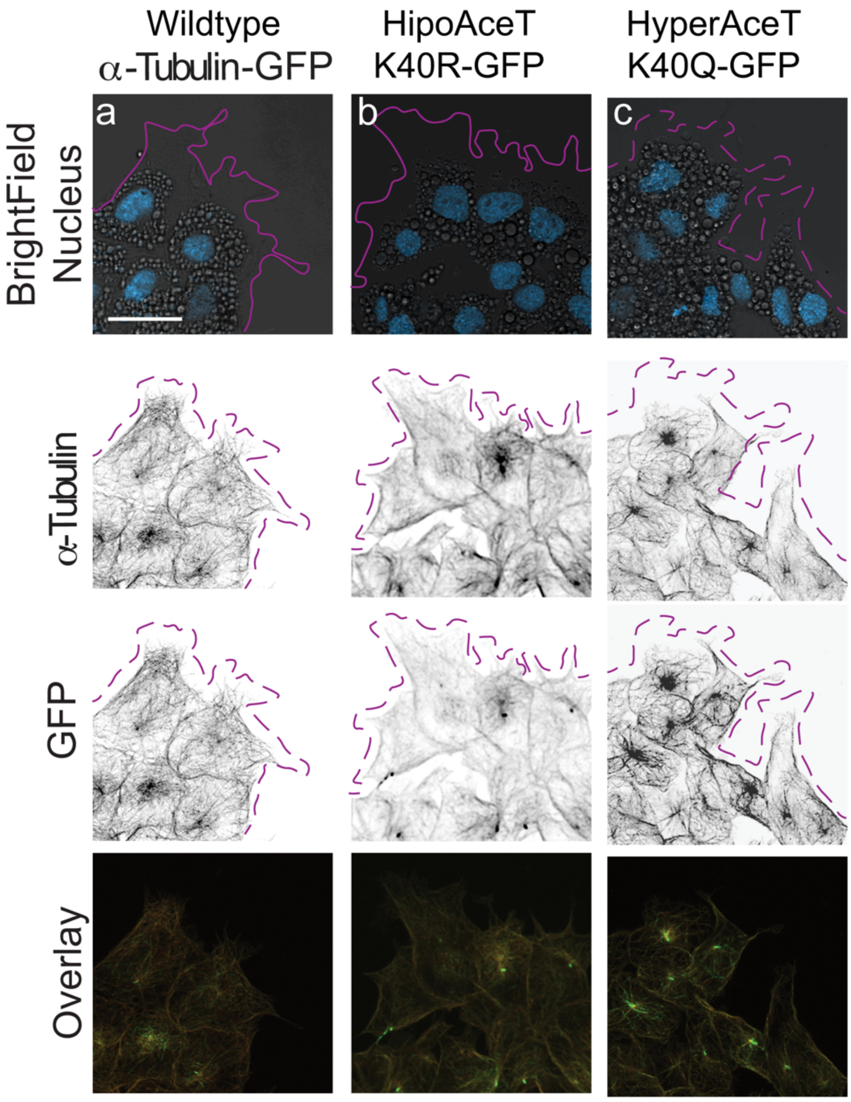


**Supplementary Figure 3. α-Tubulin constructs incorporation into endogenous microtubules.** (**a**–**c**) Incorporation of each construct is shown as indicated. Nuclei are shown in blue in the brightfield image. Note the colocalization of the endogenous α-Tubulin and with the GFP signal of each construct in the merged panels. Scale bar 20 μm. Representative examples of the data observed in our experiments.

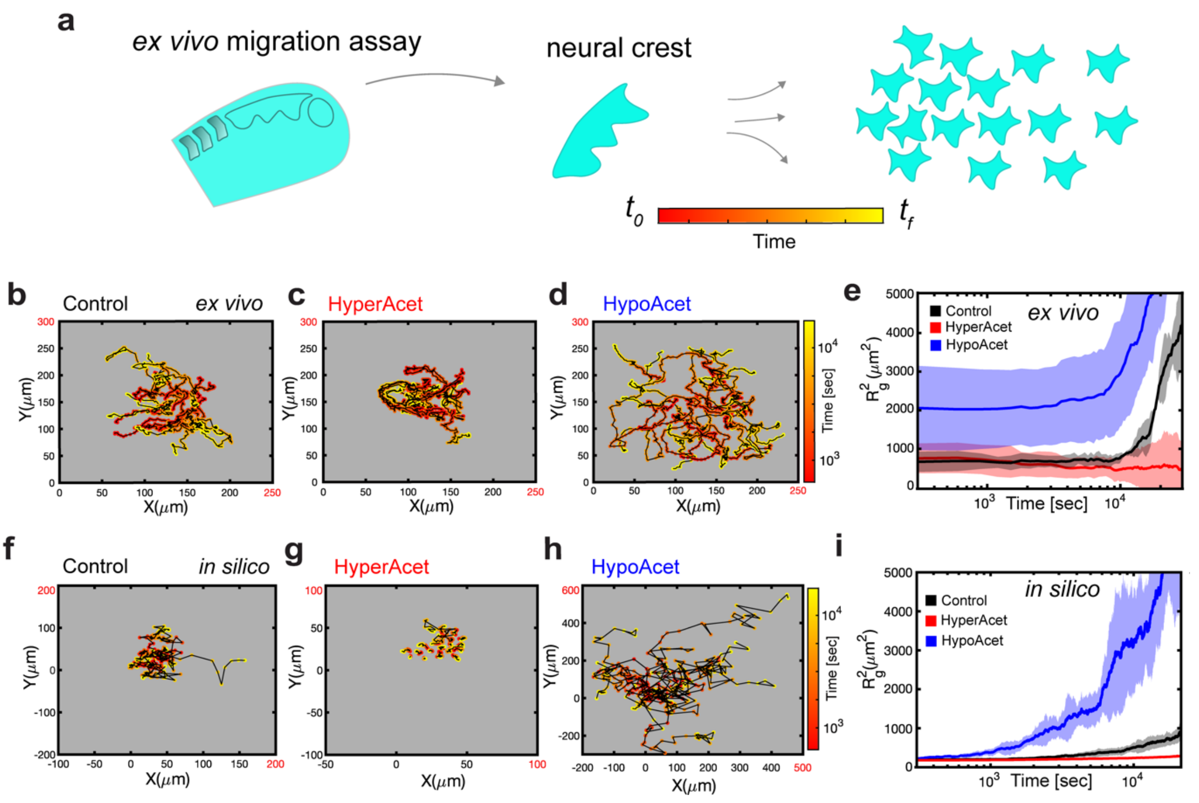


**Supplementary Figure 4. Impact of microtubule acetylation in cell migration *ex vivo* and *in silico*.** (**a**) Schematic depicts our dispersion assay (detailed in Methods). (**b–d**) *ex vivo* results for the behaviour of control, hyperacetylated and hypoacetylated neural crest cells and clusters migrating in a stiff substrate, conditions as indicated. (**b–d**) Cell tracks depicting individual cell trajectories; (**e**) $R_g^2$ *ex vivo* calculations showing cell migration under the indicated conditions, line represents average, and shadow standard deviation (SD). (**f–i**) *In-silico* results for the predicted behaviour of controls, hyperacetylated and hypoacetylated cells and clusters plated on stiff substrates. (**f–h**) Cell tracks depicting individual cell trajectories (note the differences in the x and y-axes scales when comparing, highlighted in red); (**i**) $R_g^2$ *in silico* calculations showing cell migration under the indicated conditions, line represents average, and shadow SD. **b–d** and **f–h** are representative examples from at least 3 independent experiments or simulations. Related to Figure 02 and Supplementary Movie 01.

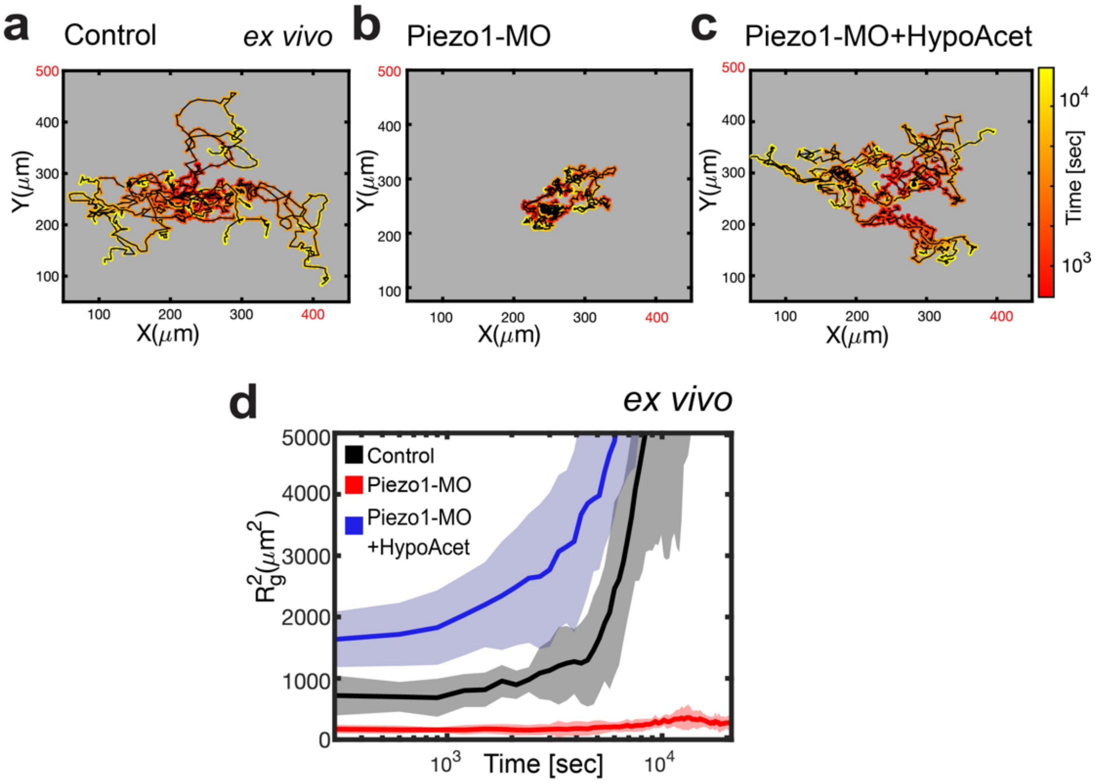


**Supplementary Figure 5. Piezo1-MO modulate cell spreading and CCM *ex vivo*.** (**a**–**d**) *ex vivo* results for the behaviour of control, Piezo1-MO, and Piezo1-MO+hypoacetylated neural crest cells and clusters when migrating on a stiff substrate; conditions as indicated in the figure. (**a**–**c**) Cell tracks depicting individual cell trajectories; (**d**) $R_g^2$ *ex vivo* calculations showing cell migration under the indicated conditions, line represents average, and shadow standard deviation (SD). **a**–**c**, representative examples of at least 3 independent experiments. Related to Figure 04 and Supplementary Movie 02.

**Supplementary Video Captions:**

**Supplementary Video 1. Microtubule acetylation modulates cell migration *ex vivo*.**
*ex-vivo* time-lapse of control, hyperacetylated and hypoacetylated neural crest cells migrating in a stiff substrate. Time-lapse setting was 1 picture every 6 min; 70 frames are shown.

**Supplementary Video 2. Piezo1 controls CCM via microtubules acetylation.**
*ex-vivo* time-lapse of control, GsMTx4, Piezo1-MO and Piezo1-MO+HypoAcet neural crest cells migrating in a stiff substrate. Time-lapse setting was 1 picture every 6 min; 70 frames are shown.

**Supplementary Table:**

| a) STRECH ACTIVATED CHANNELS | | | Reference in Mechanosensing |
|---|---|---|---|
| Protein Family | Name | TMM Normalization | References |
| PIEZO | | | |
| Piezo1 | piezo type mechanosensitive ion channel component 1 | 22.02335789 | [1] |
| Piezo2 | piezo type mechanosensitive ion channel component 2 | 10.57121179 | [1] |
| TRP Family | | | |
| *TRPA1* | transient receptor potential cation channel subfamily A member 1 | 0 | [2] |
| *TRPV1* | transient receptor potential cation channel subfamily V member 1 | 13.21401473 | [3] |
| *TRPV4* | transient receptor potential cation channel subfamily V member 4 | 6.166540208 | [4] |

**Supplementary Table 1**: **RNA-seq data from isolated neural crest cells**. While several molecules were found in our unbiassed screening, we selected just stretch activated channels that have been reported to mediate mechanosensing in other systems (see references). Next, we further filtered these candidates based on their expression levels and in their predicted role in cell migration. Since Piezo1 fulfilled these criteria, we next focused in studying the role of Piezo1 in microtubule acetylation, cell mechanics and collective cell migration.  RNA-seq protocol details can be found in (RNA-seq experimental details in **Methods**).

**Supplementary theory note on computational modelling of cell mechanical response.**

To evaluate whether cell mechanical response to microtubule (MT) acetylation facilitates collective cell migration (CCM) through cell-to-substrate stiffness mediated self-propulsion force we developed a three-dimensional active particle model using the agent-based framework. Such cell based off-lattice computational approach is known to effectively model how cell migration is impacted by cell properties such as its size, stiffness, and mechanical interaction with cell neighbours[5-9]. Individual cells are modelled as soft deformable spherical agents that interact with (i) other cells and with (ii) the substrate.

**Cell dynamics**

The net force, $\boldsymbol{F}_i$, on the $i^{th}$ cell is the vectorial sum of the forces experienced by a cell. We performed over damped (low Reynolds number[10]) dynamics without thermal noise because the viscosity is assumed to be large. Hence, the equation of motion for the $i^{th}$ cell is,

$$\dot{\boldsymbol{r}}_i = \frac{\boldsymbol{F}_i}{\gamma_i},$$

where $r_i$ is the position of the $i^{th}$ cell centre, and $\gamma_i$ is the friction coefficient. The forces experienced by a cell are described below.

**Forces**

Forces arising from cell-cell interaction, cell-substrate interaction and the active propulsion force arising from cell-to-substrate stiffness ratio are incorporated into the model. Cell-cell interaction consists of a soft repulsion term that limits spatial overlap between cells and an adhesive term accounting for cohesion between cells as mediated by cell-cell adhesion molecules. Cell-to-substrate interaction similarly accounts for a soft repulsive term that limits cell-substrate adhesion area and a cohesive term that tends to increase the adhesion area. In active particle models, a self-propulsive velocity term modelling the effect of self-generated forces in movement have been used in the context of Self Propelled Particle (SPP)[11-13] and Self Propelled Voronoi[14,15] models. While the self-propulsion term is important for modelling collective cell migration, its physical origin especially in view of the interplay between cell-substrate mechanical properties is unclear. In this context, we show that cell-to-substrate stiffness ratio is an important mediator of the self-propulsion force that cells generate to undergo migration.

Details of the force terms described above are provided below:

(i) Cell-cell interaction: The individual cells interact with other cells via short-ranged forces, consisting of elastic force (repulsion) and adhesive (attraction) force. The elastic force ($F_{ij}^{el}$) between two cells $i$ and $j$ of radii $R_i$ and $R_j$ is:

$$F_{ij}^{el} = \frac{h_{ij}^{\frac{3}{2}}}{\frac{3}{4}\left(\frac{1-\nu_i^2}{E_i} + \frac{1-\nu_j^2}{E_j}\right)\left(\frac{1}{R_i} + \frac{1}{R_j}\right)^{1/2}}$$

where $\nu_i$ and $E_i$ are the Poisson ratio and elastic modulus of the $i^{th}$ cell and $h_{ij}$ is the virtual overlap distance between the two cells. The adhesive force ($F_{ij}^{ad}$) is given by,

$$F_{ij}^{ad} = A_{ij} f^{ad} (\frac{1}{2})(c_i^{rec} c_j^{lig} + c_i^{lig} c_j^{rec})$$

where $A_{ij}$ is the overlap area between the two interacting cells and $f^{ad}$ determines the strength of the adhesive bond. We have normalized the receptor(rec) and ligand(lig) concentrations to satisfy $c_i^{rec} = c_j^{lig} = 0.9$. Cell-cell adhesion strength coefficient is fixed at $f^{ad} = 5 \times 10^{-6} \mu N/\mu m^2$ throughout the simulation.

(ii) Cell-substrate interaction: The cell-substrate elastic interaction ($F_{sub}^{el}$) is modelled based on the Hertz formalism:

$$F_{sub,i}^{el} = \frac{4}{3} \frac{R_i^{\frac{1}{2}} * \delta^{\frac{3}{2}}}{\left(\frac{1-\nu_{sub}^2}{E_{sub}} + \frac{1-\nu_i^2}{E_i}\right)}$$

where $\nu_{sub}$ and $E_{sub}$ are the Poisson ratio and elastic modulus of the substrate and $\delta$ the indentation of the cell into the substrate.

The cell-substrate adhesive interaction is given by:

$$F_{sub,i}^{ad} = A_{sub,i} f_{sub}^{ad} (\frac{1}{2})(c_{sub}^{rec} c_i^{lig} + c_{sub}^{lig} c_i^{rec})$$

where $A_{sub,i}$ is the overlap area between the a cell and the substrate and $f_{sub}^{ad}$ determines the strength of the cell-substrate adhesive bond. The substrate (sub) receptor (rec) and ligand(lig) concentrations are normalized to satisfy $c_{sub}^{rec} = c_{sub}^{lig} = 0.9$. Cell-substrate adhesion strength coefficient is set at $f_{sub}^{ad} = 9.25 \times 10^{-6} \mu N/\mu m^2$ for control cells, $f_{sub}^{ad} = 9.5 \times 10^{-6} \mu N/\mu m^2$ for hypoacetylated cells and $f_{sub}^{ad} = 9.0 \times 10^{-6} \mu N/\mu m^2$ for hyperacetylated cells. We assume that the adhesion co-efficients, receptor and ligand concentrations are constant as a function of time. As we are interested in the long-time limit of collective cell migratory behaviours (over 8 hrs), we work under the assumption that the short time fluctuations in these parameters are coarse-grained to constant values.

In addition to the mechanical interaction (elastic and adhesive forces) experienced by a cell, we incorporate a self-propulsion force $F_i^p$ that depends on the ratio of the cell-to-substrate stiffness:

$$F_i^p = T_p \left(\frac{E_{sub}}{E_i}\right)^{\frac{7}{4}} \delta \, \widehat{\boldsymbol{p}}_i$$

where $T_p$ is a propulsion force coefficient with units of tension (which we set to unity - $1\ \mu N/\mu m$), $\delta$ the cell indentation into the substrate (as defined above) and the polarity vector $\widehat{\boldsymbol{p}_i}$ specifying the direction along which the propulsion force acts. The polarity vector is assigned randomly along $\widehat{\boldsymbol{p}_i} = (\sin(\phi)\cos(\theta), \sin(\phi)\sin(\theta), 0)$ where the polar angle $\phi$ is randomly chosen in the interval $[0, \frac{\pi}{2}]$ and the azimuthal angle $\theta$ picked randomly in the interval $[0,2\pi]$. We assume that the polarity vector is not correlated in time and changes randomly with time. The out of plane component of $\widehat{\boldsymbol{p}_i}$ is set to zero to ensure that cell remains in contact with the substrate. The exponent $7/4$ gives good fit to experimentally observed time dependent cell migratory behaviours as quantified by cell spreading vs time. *Slightly modifying the exponent to smaller and larger values lead to no change in the conclusions that we report here.* In the context of purely two-dimensional motion similar self-propulsion forces have been postulated[16,17].

Friction coefficient: There are two contributions to the friction co-efficient $\gamma_i = 6\pi\eta R_i + \gamma^{max}\Sigma_{j\ in\ NN(i)}(A_{ij}\frac{1}{2}\left(1 + \frac{\vec{F}_i.\vec{n}_{ij}}{|\vec{F}_i|}\right) \times \frac{1}{2}\left(c_i^{rec}\ c_j^{lig} + c_j^{rec}c_i^{lig}\right))$. The first term is the Stokes relation $(6\pi\eta R_i)$ which models the friction with the substrate. The second friction term takes into account adhesive friction depending on cell-to-cell contact surface area $(A_{ij})$, and receptor(ligand) concentrations $(c_i^{rec}(c_i^{lig}))$. The summation is over cell nearest neighbours $NN(i)$. For any cell $i$, an array with distances from all other cells to cell $i$ is created. By calculating $R_i + R_j - |\vec{r_i} - \vec{r_j}|$ and sorting for cells $j$ satisfying $R_i + R_j - |\vec{r_i} - \vec{r_j}| > 0$ (necessary for any cell $j$ to be in contact with cell $i$) we identify the nearest neighbors.

**Simulation Details**

In each simulation, we start with placing 20 cells in a three-dimensional (3D) domain of size $X\ x\ Yx\ Z = 35\mu m\ x\ 30\mu m\ x\ 15\mu m$. The $X, Y$ positions are picked from a uniform random distribution with the bounds specified above. The margins of the $X, Y$ domain can expand (free boundary) while the z-position of the cells are constrained to be on a fixed plane at $Z = 10$. In the initial 10 steps, we allow the cells to grow in size, divide or undergo death process to facilitate randomizing the positions between simulation runs. The details of these cell processes are described in our earlier works[6]. We do not allow cells to grow in size, divide or undergo death for the rest of the simulation for a total of 3000 steps based on which we compare simulation results to experiments as we do not observe cell division, death etc during the experimental time frame. We use this scheme to randomize the initial conditions. The simulations are repeated at least 3 times per condition to ensure that initial conditions do not affect our conclusions. The time scale is assigned to 10 seconds per step (arbitrary units) to match the experimentally observed time scale of cell spreading. Codes are implemented in MATLAB.

In the simulation we model different cell stiffnesses corresponding to different levels of microtubule (MT) acetylation, keeping the substrate stiffness fixed at $E_{sub} = 100Pa$.

| **Acetylation levels** | **Cell Stiffness (Pa)** |
|---|---|
| Hypo | 75 |
| Control | 150 |
| Hyper | 400 |

For soft subtrates, the substrate stiffness is reduced to $E_{sub} = 50Pa$. The extent of cell spreading in the simulation is quantified using radius of gyration squared $R_g^2$ as discussed in the Main Text.
